# A refined *Saccharomyces cerevisiae* reference transcriptome from Direct RNA Sequencing, with a reusable pipeline for UTR annotation updates

**DOI:** 10.64898/2026.06.30.735557

**Authors:** Orlane Rossini, Alice Cleynen, Nikolay Shirokikh

**Author notes:** Corresponding authors: Alice Cleynen, Nikolay Shirokikh.

## Abstract

Untranslated regions (UTRs) flanking the coding sequence govern mRNA translation, localisation, stability, and decay, making accurate UTR boundaries essential for quantitative RNA sequencing and the study of post-transcriptional control in *Saccharomyces cerevisiae* and beyond. Reference transcriptomes built from short-read sequencing have been invaluable to the yeast community, yet in a genome as gene-dense as that of *S. cerevisiae*, short reads frequently cannot be assigned to a single transcript of origin, leaving roughly one quarter of transcripts without a confidently defined UTR. Here we use Oxford Nanopore Direct RNA Sequencing (DRS), in which each full-length polyadenylated molecule is read end to end, to resolve this ambiguity and deliver two complementary resources. First, an updated, ready-to-use *S. cerevisiae* S288C reference: change-point segmentation of per-gene DRS coverage defined boundaries for 5,416 of the 6,695 annotated genes, and a merge-max rule retaining the longer UTR from each source ensures no gene loses existing annotation. The result adds previously absent UTRs to 927 (5′) and 896 (3′) genes and extends 29.4% of 5′ and 26.1% of 3′ boundaries among comparable genes. Second, the complete, documented pipeline so that any laboratory can rebuild or update a transcriptome from its own DRS data. Validation on two independent datasets shows improved mapping rates, reduced soft-clipping, and metagene profiles consistent with genuine transcript signal.

## Introduction

Accurate gene expression measurement by RNA sequencing relies on a reference transcript model that correctly defines the boundaries of each expressed region of the DNA. Untranslated regions, the stretches of mRNA upstream (5′ UTR) and downstream (3′ UTR) of the coding sequence, are not merely spacers: they are the primary sites of post-transcriptional control, harbouring upstream open reading frames (uORFs) that modulate ribosome recruitment and translational efficiency, binding sites for RNA-binding proteins, and sequence elements governing mRNA stability, localisation, and decay (Archer et al., 2016; Hentze et al., 2018; Hinnebusch et al., 2016; Ingolia et al., 2009; Janapala et al., 2019; Shirokikh & Preiss, 2018). When UTR boundaries are under-specified in the reference (either too short or absent altogether) reads originating in the untranslated portion of the transcript fail to map, quantification estimates are biased toward the coding region alone, and any downstream biological inference built on those estimates inherits this systematic error. The practical consequences compound in proportion to the fraction of a gene’s signal that falls in its UTRs, which for many yeast transcripts is substantial.

Much of the *S. cerevisiae* UTR annotation in routine use today traces back to foundational short-read transcriptome studies, most influentially that of Nagalakshmi et al. (Nagalakshmi et al., 2008), which mapped the transcribed regions of the yeast genome at nucleotide resolution and revealed a transcriptome considerably more extensive than the annotated coding sequences alone. These resources remain widely used and are well validated for coding-region quantification. Their UTR coverage, however, is bounded not by the care of any individual study but by an intrinsic property of short-read data: a short cDNA read is a fragment and carries no information about which full-length transcript it originated from. In the compact *S. cerevisiae* genome, where adjacent genes are often separated by only a few hundred nucleotides (Goffeau et al., 1996), reads from neighbouring transcripts (whether through transcriptional read-through, alternative polyadenylation at a proximal site, or simple gene overlap) are difficult to distinguish from reads genuinely derived from the UTR of the focal gene (Tian & Manley, 2017). Boundaries must therefore be inferred from aggregate coverage patterns, and only the most unambiguous UTRs, well separated from any neighbour, can be called with confidence. As a consequence a substantial fraction of genes carry no defined UTR (approximately one quarter of protein-coding genes in current short-read-derived annotations lack a 5′ or 3′ boundary call), and the boundaries that are annotated tend toward conservative, truncated estimates.

Direct RNA Sequencing (DRS) by Oxford Nanopore Technologies (Garalde et al., 2018) resolves this limitation at the level of individual molecules. DRS captures native polyadenylated RNA without reverse transcription or fragmentation, producing reads that each represent a single, intact RNA molecule anchored at the 3′ poly-A tail. Critically, the near full-length nature of each read removes the transcript-of-origin ambiguity that confounds short-read boundary calls: a read spanning the complete extent of a transcript cannot be a contaminating fragment from an adjacent gene’s coding sequence, because it would need to originate from a polyadenylated molecule with a poly(A) tail at exactly the right genomic position. The 3′ boundary is therefore defined with high precision and specificity on every informative read. The 5′ boundary is also accessible on reads that traverse the full transcript length, though resolution here depends additionally on read completeness: reads that do not reach the transcription start site under-report 5′UTR length, a sensitivity constraint rather than an ambiguity about transcript identity. A further, systematic feature of DRS is a short blind region of approximately 14 nucleotides at the extreme 5′ terminus that is not captured in the read; because this offset is small and constant, it can be accounted for when 5′ boundaries are assigned rather than treated as a source of error.

Prior long-read studies in yeast have begun to exploit this advantage (Jenjaroenpun et al., 2018). Al Kadi et al. applied Oxford Nanopore sequencing of full-length cDNA to annotate transcript boundaries across the *S. cerevisiae* genome (the UNAGI resource), reporting improved UTR coordinates alongside novel transcripts and isoforms (Al Kadi et al., 2020). The present work differs from this and related efforts in three respects. First, we use Direct RNA Sequencing of native RNA rather than cDNA sequencing, avoiding the reverse transcription and amplification steps that can introduce length-dependent bias and displace apparent 5′ ends. Second, UTR boundary characterisation relative to the community-standard Nagalakshmi et al. reference is the primary aim, allowing a focused and exhaustively validated treatment of UTR length rather than a general transcript reconstruction. Third, we produce an explicit merge against the Nagalakshmi et al. annotation at every gene boundary, retaining the longer UTR from either source, that is formally verified to be no more restrictive than the established reference at any locus. The result is a transcript FASTA that can serve as a drop-in replacement for the standard short-read reference in any existing yeast RNA-seq pipeline, without requiring users to evaluate whether the substitution introduces regressions.

Here we present an updated *S. cerevisiae* transcript reference derived from DRS data generated in (Horvath et al., 2024) and extended by change-point segmentation of per-gene coverage profiles. We validate the reference against two independent datasets, an independent long-read DRS experiment (Fonzino et al., 2026) and the short-read Illumina data of (Al Kadi et al., 2020), across four complementary measures: read mapping rate, multi-mapping rate, antisense alignment rate, and soft-clip burden. We further validate boundary placement via metagene coverage profiles in a shared ORF-relative coordinate system, demonstrating that our extended boundaries capture genuine transcript signal rather than artefactual sequence inclusion. Together these constitute two complementary deliverables: an updated, ready-to-use *S. cerevisiae* S288C reference (transcript FASTA and GFF3 annotation) and a complete, reproducible pipeline for generating or extending such a reference from new DRS data.

## Results

### Coverage segmentation recovers gene-specific UTR boundaries from DRS data

To define transcript boundaries directly from sequencing data, we applied change-point segmentation to per-gene DRS coverage profiles spanning each annotated ORF and its 1,000 nt flanking regions using a negative binomial model (see **Methods**). Because read coverage in this window is highly autocorrelated, the raw output of Segmentor3IsBack is substantially over-segmented: a typical gene yields on the order of dozens of candidate breakpoints, the majority of which reflect local coverage fluctuation rather than genuine transcript structure. Filtering these candidates to retain only transitions where the mean coverage difference across the breakpoint exceeds 10% of the gene’s maximum transcript count collapses this redundancy into a small number of biologically interpretable segments, typically corresponding to the 5′UTR onset, the expressed ORF, and the 3′UTR decay (**Fig. 1A**).

**Figure 1.**
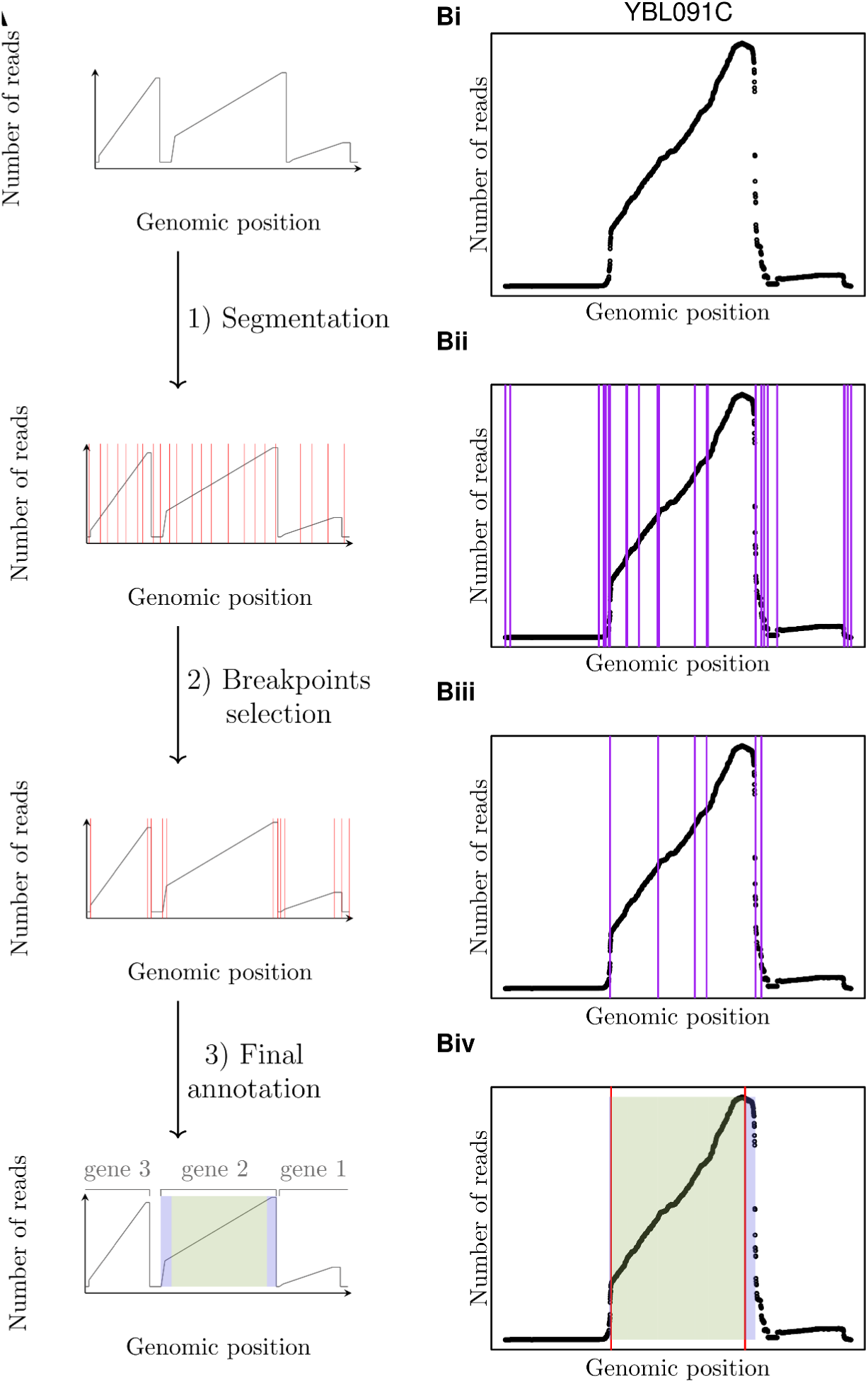
Overview of the Direct RNA Sequencing (DRS)-based UTR annotation pipeline. **(A)** Schematic illustrations of each step; **(Bi-Biv)** corresponding output applied to a representative genomic locus (YBL091C). The y-axis in all panels represents the number of reads mapped at each position; the x-axis represents genomic position. **Input (top row).** The raw DRS read-depth profile is computed across a genomic window. Because DRS reads are captured at the poly(A) tail and extend toward the 5′ end with variable length, the cumulative coverage profile rises progressively from the 3′ end of each gene and then drops sharply at the 5′ boundary, producing a characteristic staircase signal when multiple genes are present in the window. **Step 1, Segmentation.** The coverage signal is partitioned by a change-point detection algorithm that identifies positions where the slope of read depth changes significantly. All candidate breakpoints are retained at this stage (shown as vertical lines in both the schematic and the real-data example), producing an over-segmented representation of the locus. **Step 2, Breakpoint selection.** A filtering step evaluates each candidate breakpoint and retains only those supported by a sufficient number of reads and a statistically meaningful change in coverage slope. Spurious or low-confidence breakpoints are discarded, leaving a parsimonious set that corresponds to genuine transcript boundaries. **Step 3, Final annotation.** The selected breakpoints define the 5′ and 3′ UTR boundaries of each gene in the window. Genes are assigned to their respective coverage segments (illustrated with distinct coloured regions in the schematic and a shaded region in the real-data example), yielding a strand-aware UTR annotation for each locus. These per-locus annotations are subsequently merged with the Nagalakshmi et al. (2008) short-read reference, retaining the maximum boundary from either source at each end.

Figure 1 illustrates this process for a representative gene, YBL091C, chosen for its clean, well-separated data. Raw DRS coverage over the ORF ± 1,000 nt window (**Fig. 1Bi**) shows the expected profile: a sharp rise at the transcription start, a stable plateau across the expressed region, and a decline toward the 3′ end. The unfiltered segmentation (**Fig. 1Bii**) identifies 225 breakpoints distributed throughout the plateau, the majority spurious. After filtering (**Fig. 1Biii**), only the 7 breakpoints flanking the true expressed region remain, and the resulting boundaries (**Fig. 1Biv**) extend the 5′ UTR from the 7 nt reported by (Nagalakshmi et al., 2008) to 20 nt while leaving the 3′ UTR unchanged (352 nt), illustrating that the algorithm extends boundaries only where DRS coverage provides direct support for doing so, rather than systematically inflating every call.

A recurring complication in the compact *S. cerevisiae* genome is read-through from a neighbouring gene, which produces a secondary peak adjacent to the focal gene’s true expressed region. Because this secondary peak is separated from the focal plateau by a qualifying breakpoint, the filtering step correctly excludes it from the called UTR rather than incorporating it as additional transcript length.

Of the 6,695 genes in the shared gene model, 5,416 (80.9%) passed the minimum expression threshold of 20 reads at their position of maximum coverage and were carried forward for segmentation. The resulting non-zero UTR calls (prior to merging with Nagalakshmi et al.) cover 5,330 genes at the 5′ end and 5,343 genes at the 3′ end. After merging with the Nagalakshmi et al. annotation, the final reference provides a non-zero 5′ UTR for 5,805 genes and a non-zero 3′ UTR for 5,801 genes, with 5,743 genes annotated at both ends.

Additionally, UTR length can be influenced by environmental and physiological conditions (Lin & Li, 2012; Tuller et al., 2009). The inclusion of transcripts from both NS and S10 may therefore introduce a modest condition-dependent bias in the final annotation. To assess this we modeled log-transformed UTR lengths as a function of condition. For 5’ UTRs, we observed a statistically significant but small reduction in S10 compared to NS (β = −0.099, p = 5.7 × 10^-5^), corresponding to an average decrease of ∼9%; however, the effect explained very little variance (R^2^ ≈ 0.0016). For 3’ UTRs, no significant difference was detected (β = −0.023, p = 0.08; R^2^ ≈ 3 × 10^-4^). At the gene level, 69.2% of 5’ UTRs were longer or equal in NS compared to S10, while this proportion reached 78.7% for 3’ UTRs, indicating some directional asymmetry but without a strong shift in global magnitude. Overall, these results suggest that condition-dependent effects are detectable but small, and are unlikely to substantially impact the global properties of the merged annotation.

### The merged annotation systematically extends Nagalakshmi 2008 without ever shortening it

Because our final UTR length at each boundary is defined as the maximum of the DRS-derived estimate and the corresponding Nagalakshmi et al. value, a necessary property of a correctly implemented merge is that the final length should never fall below the Nagalakshmi value for any gene. We confirmed this directly: across the 4,792 genes with a comparable 5′UTR call in both annotations and the 4,817 genes with a comparable 3′UTR call, we observed zero instances in which the final annotation was shorter than Nagalakshmi at either boundary. This is not merely a sanity check on the implementation; it is the property that makes the resulting FASTA safe to adopt as a direct substitute for the Nagalakshmi reference in any existing pipeline, since no gene can lose annotated sequence as a result of the merge.

Within this conservative framework, DRS coverage extends the UTR boundary for a substantial fraction of genes. Relative to Nagalakshmi, the 5′UTR is extended for 29.4% of comparable genes and the 3′ UTR for 26.1%; the remainder are unchanged, indicating that the Nagalakshmi boundary already matched or exceeded what DRS coverage alone supported at that locus. Extensions are right-skewed: the median extension is 0 nt at both boundaries (since most genes are unchanged), while the mean extension among all comparable genes is 37.7 nt (5′) and 6.0 nt (3′), and the interquartile range for genes that are extended reaches up to 6 nt (5′) and 2 nt (3′) at the 75 ^th^ percentile overall, with a long tail of larger extensions (**Fig. 2A, B**).

**Figure 2.**
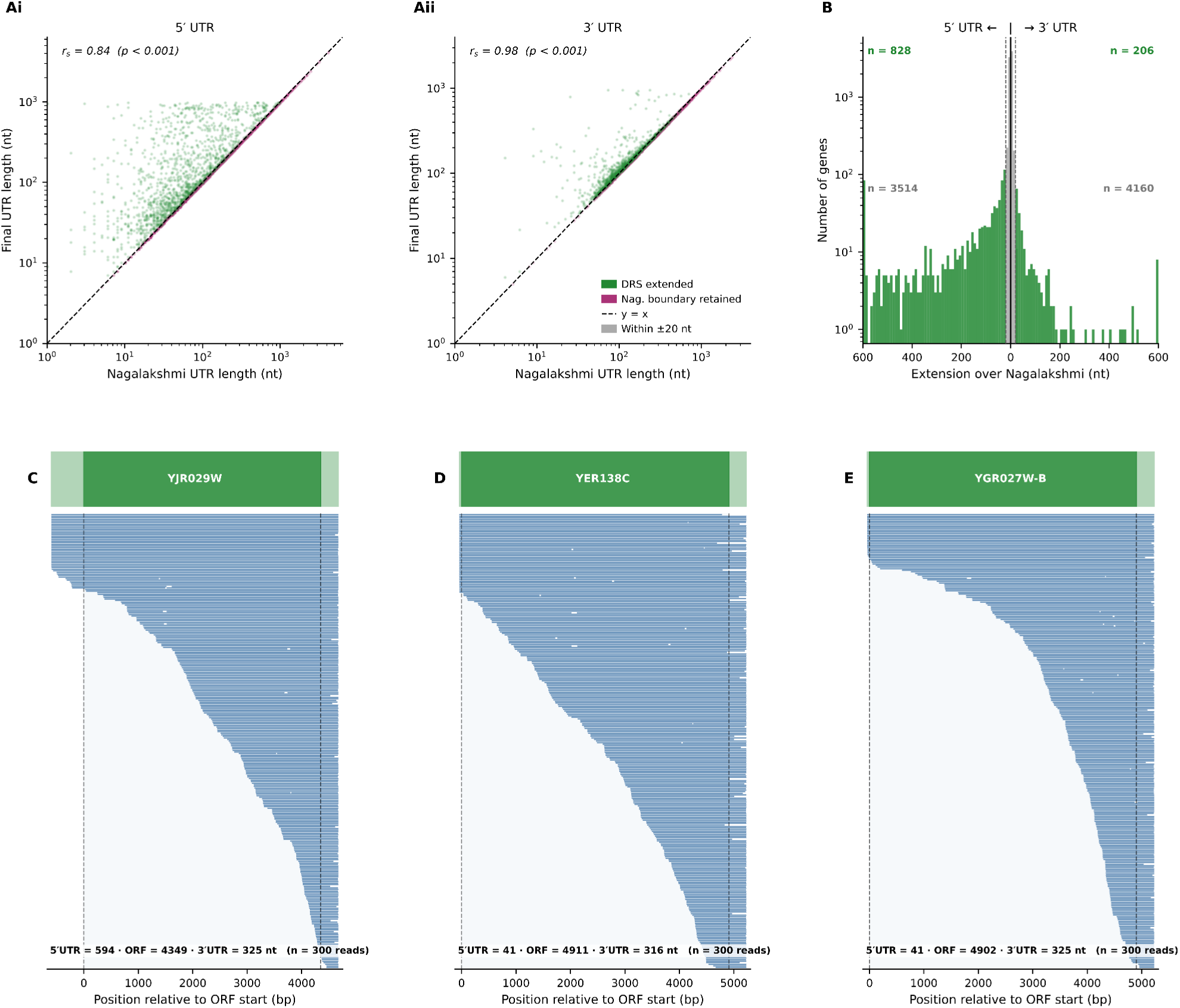
DRS-derived UTR annotation agrees with and extends the Nagalakshmi, et al. (2008) short-read reference. **(A)** Scatter plots comparing Nagalakshmi et al. UTR lengths (x-axis) with the final merged annotation (y-axis), for genes with a non-zero length reported in both sources. Left (Ai): 5′ UTR; right (Aii): 3′ UTR. Axes are log-scaled; the dashed line indicates y = x (equal lengths). Points above the identity line (green) represent genes for which Direct RNA Sequencing (DRS) extended the Nagalakshmi boundary; points on or near the line (purple) represent genes where the Nagalakshmi boundary was retained as the merge maximum. Spearman correlation coefficients (ρ) are shown. **(B)** Histograms of the difference in UTR length between the pre-merge DRS annotation and the Nagalakshmi reference (DRS − Nagalakshmi, nt) for genes with non-zero calls in both sources. Left: 5′ UTR; right: 3′ UTR. Gene counts per category are shown in log-scale. The x-axis is capped at ±600 nt; a small proportion of genes with larger differences lie outside the displayed range. **(C–E)** Representative read pileup profiles for three genes (YJR029W, YER138C, YGR027W-B), illustrating the quality of DRS read coverage used to determine UTR boundaries. Reads are shown in blue, thin strip above each pileup: final merged annotation (dark green, ORF; light green, UTRs). Dashed vertical lines mark the annotated ORF boundaries. Read counts reflect a random subsample of up to 300 primary alignments; the characteristic 5′-end coverage decline reflects the known length-dependent signal loss of Direct RNA Sequencing.

A second, distinct contribution of this resource is the recovery of UTR boundaries for genes that Nagalakshmi et al. could not annotate at all. Of the 6,607 genes in the shared gene set, Nagalakshmi reports no 5′ UTR for 1,815 genes and no 3′ UTR for 1,790 genes. DRS coverage provided a usable boundary for 927 of these genes at the 5′ end and 896 at the 3′ end. This resource directly fills roughly half of the annotation gaps left by the original short-read study. Representative examples of high-coverage genes such as YJR029W, YER138C, YGR027W-B, where DRS show read counts at the position of maximum coverage exceed 1,000, leaving little ambiguity that the called region is genuinely transcribed despite the absence of any Nagalakshmi annotation (**Fig. 2C-E**).

Where both annotations report a UTR, the two are in strong overall agreement. Because UTR length distributions are right-skewed, we report Spearman rank correlation as the primary summary statistic: ρ = 0.860 (5′ UTR, n = 4,792) and ρ = 0.979 (3′ UTR, n = 4,817); Pearson correlation gives consistent results (r = 0.922 and r = 0.984, respectively). The markedly tighter agreement at the 3′ end is consistent with the higher confidence of 3′ boundary calls from DRS data, which is anchored directly at the poly-A tail, whereas 5′ UTR length additionally depends on read completeness through the full length of the transcript (see Discussion). Restricting to genes with a real DRS-derived call on both sides (i.e. before the longest-UTR merge step), the raw DRS boundary was in fact shorter than the corresponding Nagalakshmi annotation for the majority of genes: 66.7% at the 5′ end and 69.7% at the 3′ end. To exclude differences plausibly attributable to noise in either annotation, we applied a 20 nt minimum-difference threshold; even under this conservative criterion, the DRS-derived boundary was meaningfully shorter than Nagalakshmi for 38.5% of comparable genes at the 5′ end (median shortfall 26 nt where shorter) and 52.0% at the 3′ end (median shortfall 62 nt where shorter; **Fig. 3** summaries the main differences). In every one of these cases, the final merged annotation retains the longer Nagalakshmi value rather than the shorter DRS-derived call, confirming that the longest-UTR rule performs substantial, genuine gene-by-gene arbitration between the two sources rather than defaulting to either one.

**Figure 3.**
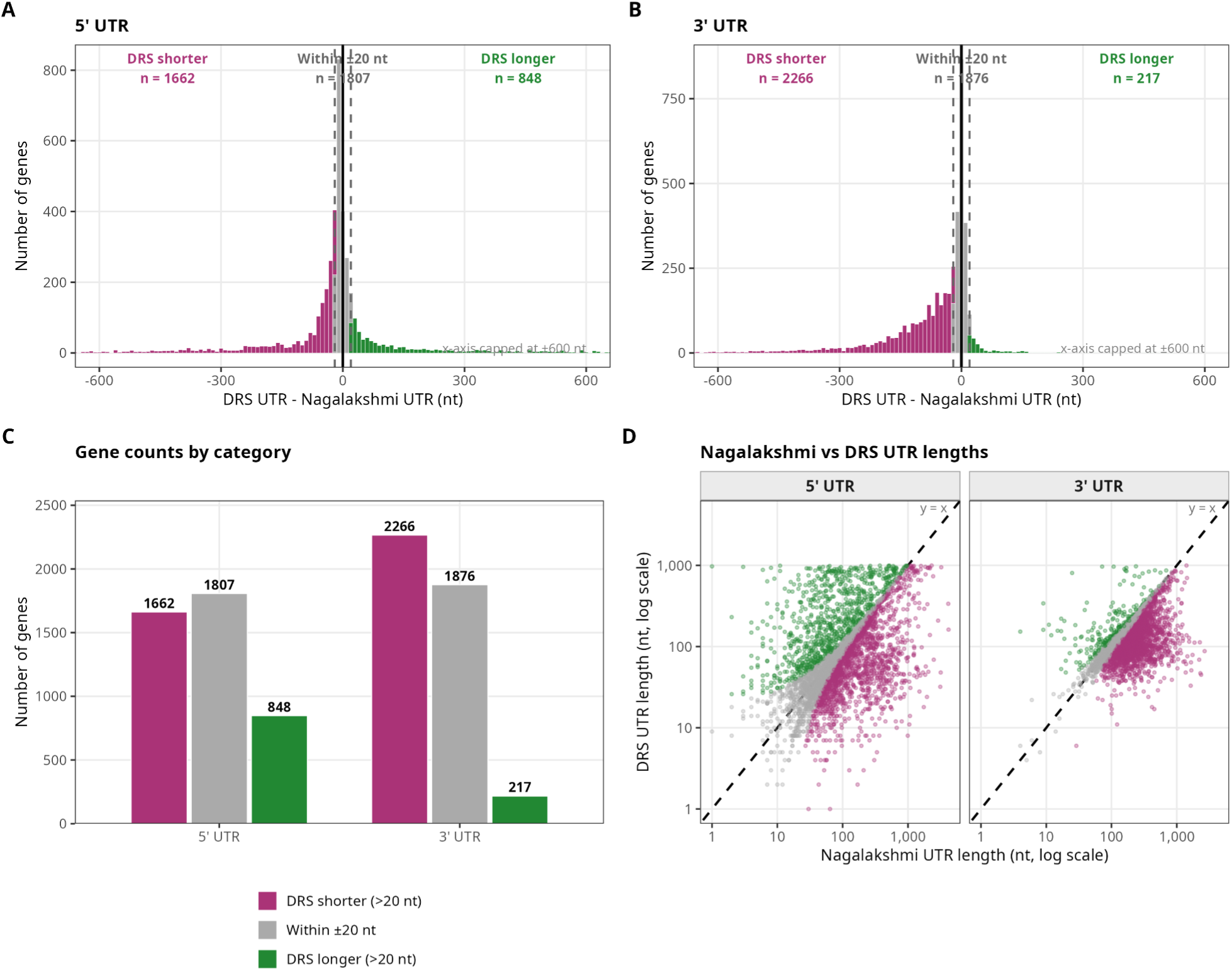
Genome-wide comparison of pre-merge DRS-derived UTR lengths against the Nagalakshmi, et al. (2008) short-read reference. Analysis restricted to genes where both the DRS annotation and Nagalakshmi et al. report a non-zero UTR length at each end. Genes are classified into three categories based on a ±20 nt threshold (chosen to exclude differences attributable to measurement noise): DRS shorter (purple, difference ≤20 nt), within ±20 nt (grey), and DRS longer (green, difference > +20 nt). **(A, B)** Histograms of the difference in UTR length (DRS − Nagalakshmi, nt) for the 5′ UTR (A) and 3′ UTR (B). The x-axis is capped at ±600 nt; a small fraction of genes with more extreme differences fall outside the displayed range. **(C)** Grouped bar chart summarising gene counts in each category for both UTR ends. At the 5′ end, DRS calls are shorter than Nagalakshmi for a larger fraction of genes (n=1,662 shorter vs. n=848 longer), consistent with known 5′-end signal dropout inherent to DRS. At the 3′ end, this bias is more pronounced (n=2,266 shorter vs. n=217 longer). **(D)** Scatter plots (log–log scale) of Nagalakshmi UTR length (x-axis) vs. DRS UTR length (y-axis) for the 5′ (left) and 3′ (right) UTR, coloured by category. The dashed line indicates y = x (equal lengths). The overall correlation confirms that the two annotation sources broadly agree on relative UTR length, with the merge procedure preserving the maximum boundary from either source.

As a fully independent validation, we compared the final annotation against the TIF-seq boundaries (Pelechano et al., 2013), an orthogonal method that maps both transcript ends directly and was not used in constructing the reference. Across the genes with a covering major transcript isoform (n = 5,360 at the 5′ end and n = 5,413 at the 3′ end), the two annotations were positively correlated (Spearman ρ = 0.48 and ρ = 0.55, respectively). Critically, the final annotation under-called the independent TIF-seq boundary by more than 20 nt for only 4.2% of genes at the 5′ end (225 of 5,360) and 4.9% at the 3′ end (264 of 5,413); the remaining genes were either concordant within 20 nt or carried a longer final boundary. This confirms that, even against an orthogonal method, the reference rarely truncates transcript ends, while the predominance of longer-final cases is consistent with the conservative, longest-UTR design (**Fig. 4**).

**Figure 4.**
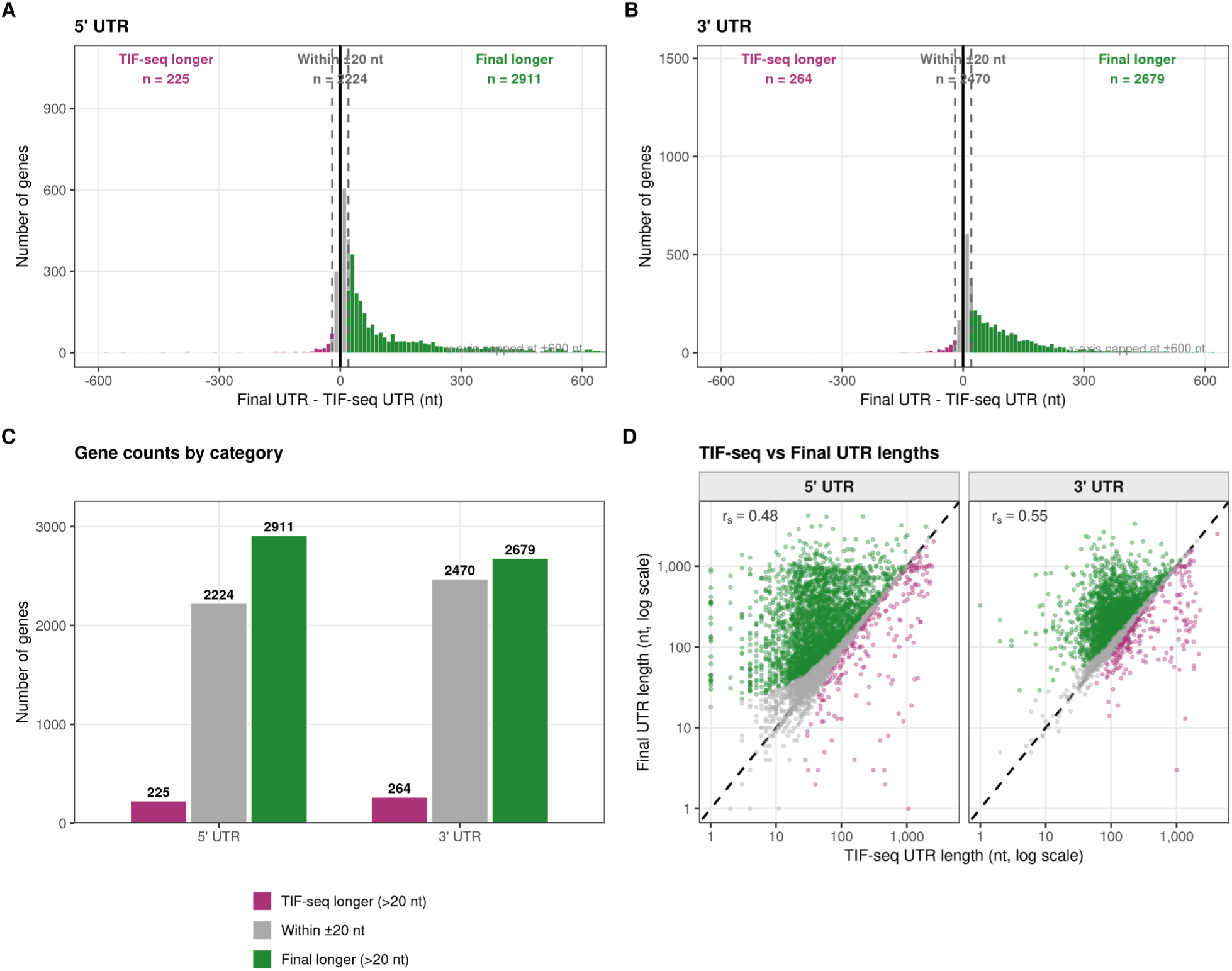
Genome-wide comparison of the final merged UTR annotation against independent TIF-seq transcript-isoform boundaries (Pelechano *et al*., 2013). For each gene, the final annotation was compared with the major covering transcript isoform (mTIF) derived from TIF-seq, an orthogonal method that maps the 5′ and 3′ ends of individual transcripts simultaneously and is fully independent of the data used to construct the reference. (A, B) Distribution of the signed difference (final UTR − TIF-seq UTR) at the 5′ (A) and 3′ (B) ends; positive values indicate a longer final boundary. The solid line marks no difference and the dashed lines the ±20 nt threshold; the x-axis is capped at ±600 nt. (C) Genes classified by the ±20 nt threshold as TIF-seq longer, concordant (within ±20 nt), or final longer, shown separately for each end. (D) Final versus TIF-seq UTR length on log–log axes, coloured by category, with Spearman rank correlation (5′ UTR ρ = 0.48, n = 5,360; 3′ UTR ρ = 0.55, n = 5,413). Across both ends the final annotation under-calls the TIF-seq boundary for fewer than 5% of genes (4.2% at the 5′ end, 4.9% at the 3′ end), confirming that the reference rarely truncates transcript ends relative to an independent benchmark.

### The merged reference improves read mapping relative to naive alternatives, on both long- and short-read data

To evaluate the practical effect of UTR definition on downstream alignment, we mapped both an independent DRS data (Fonzino et al., 2026) and an independent short-read dataset (Al Kadi et al., 2020) to five references built from the same underlying gene models: the full genome, an ORF-only transcriptome, a transcriptome with a fixed ±1,000 nt flank around each ORF, the Nagalakshmi et al. annotation alone, and our final merged annotation ("our ref"). Because the fixed-flank and ORF-only references share no biological basis for their boundaries, they serve as naive baselines against which to judge whether UTR-aware references offer a genuine improvement, and whether that improvement extends beyond the data used to construct it.

For the DRS data (3,423,258 primary reads), mapping to our reference recovered a 94.14% mapping rate, closely matching the genome (95.21%) and Nagalakshmi (93.37%) references and substantially exceeding the ORF-only transcriptome (85.38%), which by construction provides no homology target for reads originating in untranslated regions (**Fig.5A**). Critically, this gain in mapping rate was not accompanied by the loss of specificity seen in the naive ±1,000 nt flank reference, which achieved a similar mapping rate (94.36%) but at a multi-mapping rate of 84.21%, or more than five times that of our reference (15.31%), reflecting the fact that fixed flanks routinely capture sequence from neighbouring genes in the compact yeast genome (**Fig. 5B**). Antisense mapping was lowest for our reference of all five (5.96%), below both Nagalakshmi (6.62%) and ORF-only (8.59%), indicating that the merged boundaries are not simply longer but remain well localised to genuine sense-strand transcript sequence rather than incidentally capturing adjacent or antisense signal (**Fig. 5C**).

**Figure 5.**
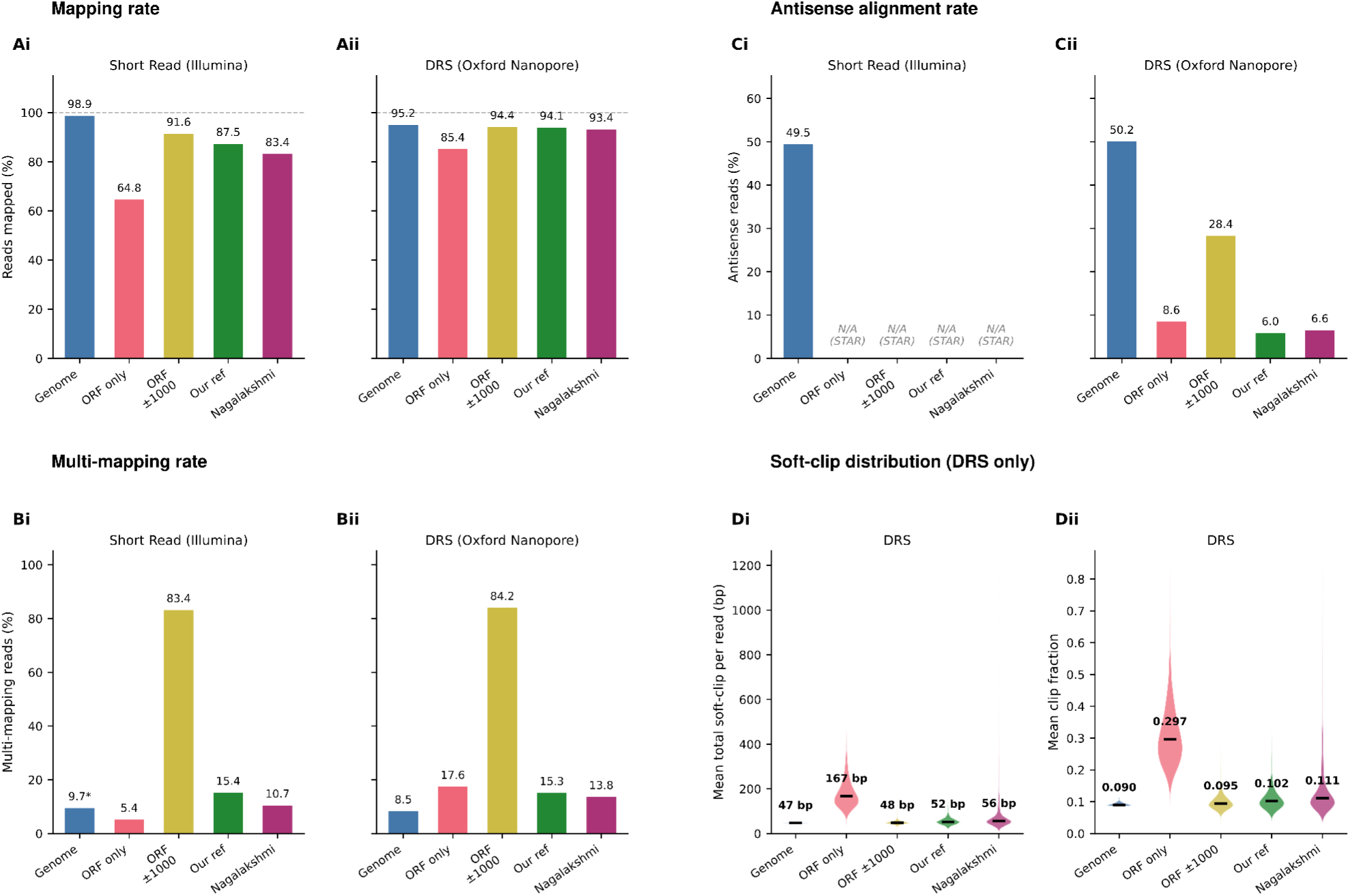
Alignment quality metrics across five reference transcriptomes for short-read (Illumina) and Direct RNA Sequencing (DRS, Oxford Nanopore) data. **(A)** Mapping rate for short-read (Ai) and DRS (Aii) data. Values represent the percentage of input reads that map as primary alignments to each reference. The denominator is the raw FASTQ read count for short reads and the total reads in the BAM (including unmapped) for DRS. The dashed line indicates 100%. **(B)** Multi-mapping rate for short-read (Bi) and DRS (Bii) data. For the genome reference aligned with bwa-mem2 (asterisk), multi-mapping is estimated as the fraction of primary alignments with MAPQ = 0, since bwa-mem2 does not report secondary alignments (flag 0x100). For STAR-aligned transcriptome references, multi-mapping is defined as the fraction of reads carrying an NH tag ≥ 2. For DRS (minimap2), multi-mapping is defined as reads with at least one secondary alignment. The exceptionally high rate for ORF±1000 reflects the dense transcript overlap inherent to flanking-extended references in the compact yeast genome. **(C)** Antisense alignment rate for short-read (Ci) and DRS (Cii) data (primary mapped reads only). STAR suppresses antisense alignments for transcriptome references by default; these references are therefore marked N/A. **(D)** Per-transcript soft-clip distribution for DRS reads mapped to each reference (DRS only). Di shows the mean total soft-clipped bases per read (bp); Dii shows the mean soft-clip fraction (clipped bases / read length). Violin bodies show the distribution across transcripts; horizontal bars indicate medians (annotated). ORF-only references show markedly higher soft-clipping, consistent with reads regularly extending beyond the reference boundaries into un-annotated UTR sequence. Our reference and Nagalakshmi show similarly low soft-clip fractions, comparable to the full-genome reference.

A complementary line of evidence comes from soft-clip length, which measures how much of each aligned read falls outside the reference sequence rather than whether the read maps at all. Across the 5,861 transcripts annotated in both our reference and Nagalakshmi, median total soft-clip length (5′ + 3′ combined) was lower for our reference at every position in the distribution we examined: 52.3 nt versus 55.9 nt overall, and lower at each end individually (5′: 3.1 vs 3.6 nt; 3′: 47.7 vs 50.0 nt), with a mean per-read reduction of

17.7 nt when compared transcript-by-transcript. Because soft-clipped bases represent sequence the aligner could not place within the reference, this indicates that our boundaries capture more of the true transcript within the annotated sequence itself, rather than leaving it to be clipped at alignment time. The ORF-only reference shows by far the highest clip burden of the four (median 167.3 nt), as expected for a reference that provides no UTR sequence at all; the ±1,000 nt flank reference shows the lowest absolute clip values of any reference (48.2 nt), but this reflects the generosity of a flank far larger than any real UTR rather than a more accurate boundary, and should be read alongside its substantially worse multi-mapping and antisense rates above rather than as evidence of superior boundary placement (**Fig. 5D**).

This pattern was independently reproduced on short-read data that played no role in constructing the annotation. Of 34,270,624 total short reads, 29,992,595 (87.51%) mapped to our reference, compared with 28,594,996 (83.43%) to Nagalakshmi and 22,216,695 (64.82%) to the ORF-only transcriptome, a gain of approximately 1.4 million additional mapped reads relative to Nagalakshmi alone, recovered without any change to the underlying sequencing data. As with the DRS data, the ±1,000 nt flank reference reached a nominally higher mapping rate (91.63%) only by accepting a correspondingly inflated multi-mapping rate (83.37%, versus 15.39% for our reference), confirming that this naive baseline trades specificity for coverage rather than offering a genuine improvement. Together, these results show that the merged reference inherits the clean mapping behaviour of the Nagalakshmi annotation while recovering additional correctly mapped reads, on both the data used to build it and an independent dataset, and does so without the multi-mapping cost of simply widening the reference indiscriminately. **Fig. 6** shows representative read pileups from both independent datasets for direct visual comparison.

**Figure 6.**
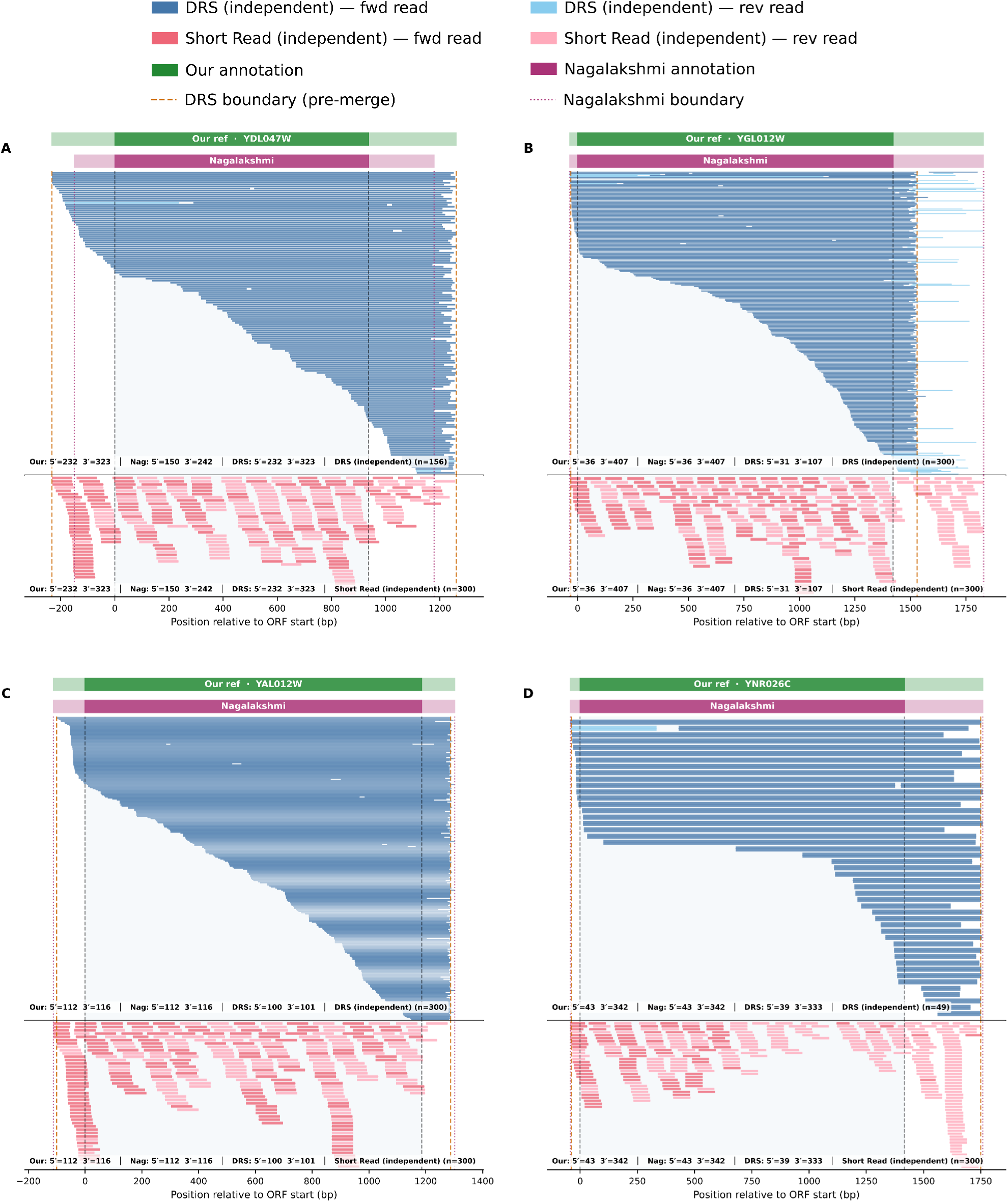
Independent validation of DRS-derived UTR boundaries using DRS and short-read (Illumina) coverage from datasets distinct from those used for annotation. Each panel shows read pileups from two independent datasets, DRS (blue, upper sub-panel) and short-read Illumina (salmon, lower sub-panel), both mapped to our merged reference annotation, plotted relative to the annotated ORF start (position 0). Two annotation tracks are shown above each pileup: the final merged annotation (green, "Our ref") and the Nagalakshmi *et al*. reference (purple). Solid black dashed vertical lines mark ORF boundaries; orange dashed lines indicate the pre-merge DRS-derived boundaries; purple dotted lines indicate the Nagalakshmi boundaries. The characteristic DRS coverage pattern (progressive 5′-end dropout) is visible in the upper sub-panels; the short-read panels show dense, uniform coverage throughout the transcript. UTR lengths for all three sources are shown bottom-left of each sub-panel. Up to 300 reads are displayed per panel. **(A) YDL047W, a discordant case.** The DRS annotation extends both UTR boundaries beyond the Nagalakshmi reference (5′: 232 *vs.* 150 nt; 3′: 323 *vs.* 242 nt). Both the independent DRS and short-read datasets confirm coverage initiating and terminating beyond the Nagalakshmi boundaries at both ends, supporting the longer DRS-derived boundaries adopted in the merge. **(B) YGL012W, a discordant case (3′ end).** The pre-merge DRS annotation yields a substantially shorter 3′ UTR than Nagalakshmi (107 *vs.* 407 nt). The independent short-read coverage extends well beyond the DRS 3′ boundary and is consistent with the Nagalakshmi estimate of 407 nt, confirming that the DRS truncation reflects 3′-end signal loss rather than a genuine biological boundary. The merged annotation correctly retains the Nagalakshmi 3′ UTR. **(C) YAL012W, a concordant case.** DRS and Nagalakshmi boundaries agree closely at both ends (5′: 100 *vs.* 112 nt; 3′: 101 *vs.* 116 nt, both within ±20 nt). Both independent datasets show consistent coverage up to the annotated boundaries, with no signal beyond them, confirming high-confidence annotation at this locus. **(D) YNR026C, a concordant case.** DRS and Nagalakshmi boundaries agree almost to the nucleotide (5′: 39 *vs.* 43 nt; 3′: 333 *vs.* 342 nt). Both independent DRS and short-read profiles terminate sharply at the merged boundaries, providing strong cross-platform support for the final annotation.

### Metagene coverage confirms that extended boundaries capture genuine transcript signal

The mapping-rate comparisons establish that more reads map to our reference overall, but not where those reads map relative to the annotated boundary. To address this directly, we computed metagene coverage profiles anchored at the transcript start (TSS) and end (TES) for all four transcriptome references, using a shared ORF-relative coordinate system so that mean read depth at a given position is directly comparable across references, and restricting to the 4,791 genes where Nagalakshmi had defined UTR bounds (**Methods**).

The ORF-only reference shows the expected null result by construction in both data types: mean read depth is exactly zero at every position upstream of the annotated start and downstream of the annotated end (DRS: 467.9 reads per base at −10 nt falling to 0 from position 0 onward at the TES; short reads: 276.3 reads per base at −10 nt falling to 0 at the same boundary), since no sequence exists there for reads to align to ( **Fig. 7**).

**Figure 7.**
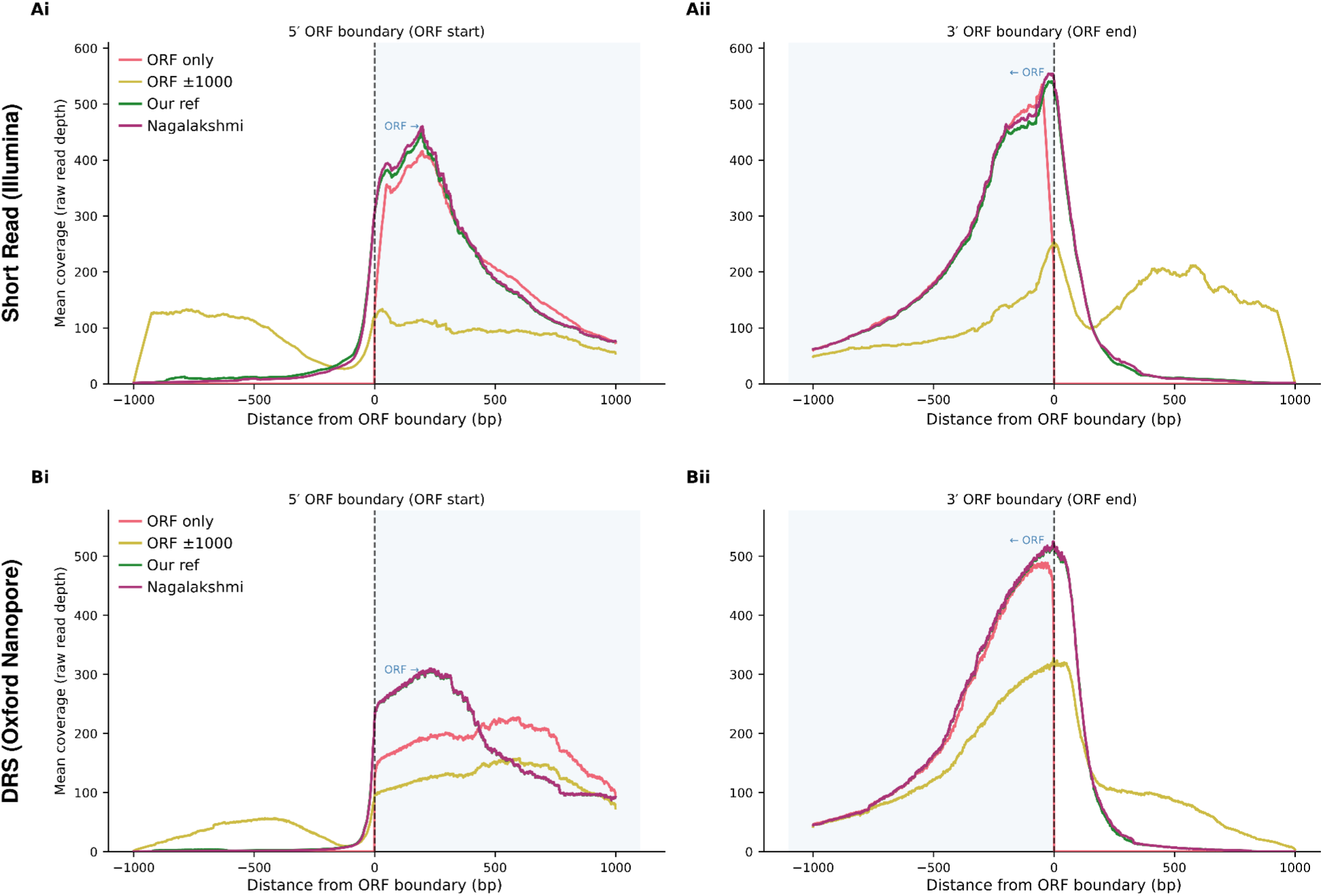
Metagene coverage profiles around ORF boundaries for four transcriptome references. Mean read depth is plotted as a function of distance from the annotated ORF boundary (position 0, dashed line) across all genes passing the coverage threshold (∼4,696–4,700 transcripts per reference, restricted to the shared gene set). The light blue shaded region indicates the ORF interior. Profiles are shown for four references: ORF-only (pink), ORF±1000 (yellow), our merged DRS annotation (green), and Nagalakshmi et al. (purple). **(A)** Short-read (Illumina) metagene profiles anchored at the 5′ ORF boundary (ORF start, Ai) and 3′ ORF boundary (ORF end, Aii). Our reference and Nagalakshmi show near-identical profiles, with coverage rising sharply at the ORF boundary and the characteristic upstream signal of reads spanning the 5′ UTR. The ORF-only reference produces a very similar intra-ORF profile but drops abruptly at the boundary due to the absence of UTR sequence. ORF±1000 shows substantially depressed coverage throughout, consistent with the high multi-mapping rate caused by the 1000 nt flanks in the compact yeast genome diluting signal across overlapping transcripts. **(B)** DRS (Oxford Nanopore) metagene profiles at the same boundaries (Bi: ORF start; Bii: ORF end). The characteristic DRS signature of reads anchored at the poly(A) tail and declining in coverage toward the 5′ end produces a gradient of coverage across the ORF, most visible in Bi where coverage increases from left to right as reads grow shorter and fewer reach the 5′ end. At the 3′ boundary (Bii), our reference and Nagalakshmi again co-localise closely, with a sharp coverage peak at position 0 reflecting accurate 3′ end annotation. ORF-only shows a comparable peak, while ORF±1000 is again substantially suppressed.

The ±1,000 nt flank reference, in contrast, does not merely show residual coverage in its fixed flank: it shows coverage that approaches, and in places exceeds, the level seen within the ORF body itself, the clearest possible signature of contamination from neighbouring genes. In the DRS data, coverage at 500 nt upstream of the TSS reaches 53.9 reads per base in the ±1,000 nt reference, compared with a typical ORF-body level of around 121 reads per base measured at 800 nt into the gene in the same reference, that is, coverage nearly halfway back to ORF levels at a position that, in our reference and in Nagalakshmi, has already decayed to background (2.2 and 1.6 reads per base respectively). At the TES, the effect is more pronounced still: coverage 500 nt downstream of the annotated end reaches 86.3 reads per base in the ±1,000 nt reference, *higher* than the 63.3 reads per base measured 800 nt before the boundary in the same reference, while our reference and Nagalakshmi have decayed to 7.1 reads per base at the same distance. The short-read data show the same pattern in even starker form: at 500 nt upstream of the TSS, ±1,000 nt coverage reaches 111.3 reads per base, exceeding the reference’s own ORF-body coverage of 82.6 reads per base measured at 800 nt into the gene, and at 500 nt downstream of the TES it reaches 198.6 reads per base against an ORF-body level of 64.2 reads per base measured at 800 nt before the boundary in the same reference, i.e. in these cases coverage in the fixed flank is, if anything, higher than coverage inside the annotated coding sequence. This pattern is only explicable by reads originating from the coding sequences of adjacent genes mapping indiscriminately into the fixed flank of the focal gene, a direct consequence of the dense, closely packed gene arrangement of the *S. cerevisiae* genome; it cannot reflect genuine UTR signal for the focal transcript; and it is precisely the artefact that boundary-aware references, by construction, are immune to.

Our reference and the Nagalakshmi reference both show the expected biological profile at each boundary in both data types: a gradual rise in mean read depth approaching the TSS (DRS: 0.57 to 227 reads per base between −1,000 nt and the annotated ORF start for our reference, 0.60 to 228 reads per base for Nagalakshmi; short reads: 1.71 to 305 reads per base for our reference, 1.76 to 307 reads per base for Nagalakshmi) and a corresponding gradual decay beyond the TES, rather than the step-function seen in ORF-only or the inflated, non-decaying tail seen in the ±1,000 nt reference. The two UTR-aware references are essentially indistinguishable at the ORF boundary itself; at positions further into the upstream UTR region (e.g. 500 nt upstream of the TSS, where our reference shows 2.2 versus 1.6 reads per base for Nagalakshmi), our annotation produces marginally higher mean depth, consistent with a subset of genes carrying extended 5′UTRs that contribute signal at those upstream positions. The reproduction of every feature of this pattern in the independent short-read dataset rules out the possibility that the extended boundaries are a DRS-specific mapping artefact.

## Discussion

### A new transcriptome reference for yeast, built for direct use

The yeast research community has lacked an updated, readily usable UTR annotation since Nagalakshmi et al. established the field-standard reference using short-read tiling arrays and early RNA-seq (Nagalakshmi et al., 2008). That resource has been remarkably durable, but it carries the structural limitations of the technology that produced it: short-read coverage declines gradually toward transcript ends, and boundary calls made from declining coverage are inherently conservative, with no annotation possible at all for genes whose expression or local genome architecture left the signal too ambiguous to call. Direct RNA Sequencing removes this limitation at its source by capturing each transcript as a single molecule from 3′ poly(A) to (where read length allows) the immediate vicinity of 5′ cap, making both UTR boundaries directly observable on the same read rather than inferred from the decay of pooled short-read coverage.

The contribution of this work is not simply a new annotation but a complete, directly substitutable transcriptome reference: a merged FASTA, derived by retaining the longer UTR call between our DRS-based segmentation and the Nagalakshmi annotation at every boundary, together with the documented and reproducible pipeline used to build it. This dual deliverable matters for two different audiences. For most users, the FASTA itself is the resource: a drop-in replacement for any existing yeast RNA-seq workflow currently built around Nagalakshmi or an ORF-only reference, requiring no reprocessing of raw data and no familiarity with the underlying segmentation method. For users with their own DRS data, or who wish to apply the same logic to a different strain, condition, or organism, the pipeline itself is reusable, and we provide it openly for that purpose.

### Improvement over the Nagalakshmi 2008 reference

Three lines of evidence in this work support the claim that the merged reference is a genuine improvement rather than a cosmetic update. First, the merge construction guarantees that no gene loses annotated UTR sequence relative to Nagalakshmi, a property we confirmed held without exception across thousands of comparable genes (**Results**). Second, where DRS coverage supports a longer call, it does so for a substantial minority of genes at each boundary, however most importantly it supplies a usable boundary for roughly half of the genes Nagalakshmi could not annotate at all, recovering information that has simply been absent from the field’s reference annotation for over a decade. Third, and most importantly for practical adoption, these extensions are not an artefact of the merge construction: independent mapping-quality and metagene analyses, performed on data not used to build the annotation, show that reads in the extended regions behave exactly as genuine transcript signal should: they map with high specificity, are not enriched for soft-clipping, and form a smooth, biologically shaped coverage profile rather than the elevated, non-decaying signal seen when a reference’s flank is defined arbitrarily rather than from data.

The asymmetry between the 5′ and 3′ results is itself informative. Agreement between our calls and Nagalakshmi’s is markedly tighter at the 3′ end (Spearman ρ = 0.979) than at the 5′ end (ρ = 0.860), and the mean extension at the 5′ end is several-fold larger than at the 3′ end. This is consistent with the underlying biology of DRS: sequencing proceeds 3′ to 5′ from the poly-A tail, so the 3′ boundary is anchored directly and with high confidence on essentially every read that maps to a transcript, while the 5′ boundary is only as reliable as the longest reads that survive intact to the start of the molecule (Workman et al., 2019). Shorter or degraded reads systematically under report 5′ UTR length, meaning our 5′ UTR calls likely still understate the true 5′ boundary for at least some genes, even after the conservative merge with Nagalakshmi. This is a meaningful caveat for any 5′-UTR-specific analysis (translation initiation, upstream ORF usage) built on this resource, and is taken up further below.

### Relationship to comparable long-read efforts in yeast

The most directly comparable prior effort is the UNAGI pipeline of (Al Kadi et al., 2020), which applied Nanopore long-read sequencing of full-length cDNA to improve transcript annotation in *S. cerevisiae*, including 5′ and 3′ UTR boundaries. The present work differs from UNAGI in scope, technology, and deliverable. In scope, UNAGI’s primary contribution is the discovery of novel transcripts and isoforms genome-wide, with UTR annotation as one component of a broader transcript-reconstruction pipeline. Here, UTR characterisation relative to an existing, near-complete gene model is the central question, allowing a more targeted and exhaustively validated treatment of UTR length specifically. In technology, the present work uses Direct RNA Sequencing, which sequences native RNA without a reverse-transcription or PCR amplification step and is therefore not subject to the strand and length biases that can accompany cDNA synthesis; UNAGI, by contrast, sequenced full-length cDNA. In deliverable, this work produces an explicit, principled merge against the existing community reference whereas UNAGI’s output is a transcript reconstruction rather than a backward-compatible drop-in replacement for the Nagalakshmi annotation specifically.

More broadly, this work sits alongside a small but growing body of long-read transcriptomics in yeast, including UNAGI and the FLAIR-based comparisons it reports (Tang et al., 2020), that collectively make the case that short-read-derived reference annotations from the 2000s era systematically understate UTR extent in compact genomes. We see this resource as complementary to, rather than a replacement for, that broader effort: where UNAGI and similar tools are best suited to discovering novel transcripts and isoforms, the present resource is purpose-built for users whose primary need is an accurate, ready-to-use UTR annotation compatible with the existing Nagalakshmi-based literature.

### The choice to take the maximum UTR length, and its limitations

The central design decision in this work to retain the longer of the DRS-derived and Nagalakshmi UTR length at each boundary, independently per gene and per side was made to prioritise backward compatibility and safety of adoption over maximal data-driven accuracy. We discuss here both the justification for this choice and its limitations.

The justification is principally practical. A new annotation that replaced Nagalakshmi outright would require every adopting laboratory to evaluate whether the new resource introduces regressions relative to whatever analyses they currently run on the established annotation; for most users, the cost of that evaluation outweighs the benefit of a few hundred nucleotides of additional UTR sequence on a subset of genes. By construction, the longest-UTR merge cannot regress any existing analysis built on Nagalakshmi coordinates: any region annotated by Nagalakshmi remains annotated, and the only possible change is the inclusion of additional sequence beyond what Nagalakshmi reported.

This choice has two related limitations. First, it is conservative by design and therefore does not correct cases in which the Nagalakshmi annotation may be too permissive at a locus; for instance where short-read coverage extended into what later, longer-read evidence would identify as the start of a neighbouring transcript. The merge logic has no mechanism to shorten a Nagalakshmi call, even where DRS evidence might support doing so; we confirmed that such cases do occur (the DRS-derived boundary was, for more than half of the genes, shorter than the corresponding Nagalakshmi call; see **Fig. 3**), and a fully data-driven re-annotation, unconstrained by backward compatibility, would in principle be free to revise such cases. We view this as an acceptable trade-off for a resource paper rather than a shortcoming to be corrected: a maximally accurate but non-backward-compatible annotation and a conservative, backward compatible one serve different purposes, and we have deliberately built the latter.

Second, and more subtly, “maximum across sources” is not the same guarantee as “correct.” Because our own DRS-derived estimate is itself taken as the maximum UTR length observed across all sequencing conditions and RNA fractions (**Methods**), and is then merged with Nagalakshmi by retaining whichever of the two is longer, the final annotation is, by construction, the longest plausible UTR call available to us from any source examined. This maximises sensitivity at the cost of an unquantified increase in false-positive UTR length for at least some genes, for example where a single sequencing condition or fraction captured an unusually long but non-representative read, or where neighbouring-gene contamination was not fully excluded by our segmentation filter despite the safeguards described in **Methods**. We have not attempted to quantify the rate of such over-calls directly, and consider this an open question for future refinement of the resource rather than one we can resolve with the present data. We note, however, that the statistical effect of condition on UTR length is small, providing empirical reassurance that the pooling of conditions does not substantially distort the global annotation.

### Outlook and applications

For the practising yeast researcher, the immediate value of this resource is quantitative: counting reads against correct UTR boundaries reduces the systematic loss of signal that occurs when transcript ends are truncated or absent, improving differential-expression and transcript-abundance estimates without any change to existing workflows. Beyond quantification, the recovered and extended UTRs open specific lines of enquiry that an ORF-only or UTR-incomplete reference cannot support: the 3′ UTRs underpin studies of alternative polyadenylation, mRNA stability (Tudek et al., 2021), and the *cis*-regulatory elements bound by RNA-binding proteins, while the 5′ UTRs (with the sensitivity caveat noted above) inform analyses of translation initiation and upstream open reading frames. Because the pipeline is released alongside the annotation, the same approach can be applied to other *S. cerevisiae* strains, to defined growth or stress conditions, or to non-conventional yeasts and other compact genomes, yielding condition- or strain-specific references where a single static annotation is inadequate.

Several extensions follow naturally from this work. The conservative, backward-compatible merge could be complemented by a fully data-driven re-annotation permitted to shorten over-extended boundaries, once the rate of such cases can be estimated against orthogonal evidence. Our comparison against transcript-isoform sequencing TIF-seq (Pelechano et al., 2013), which delimits both transcript ends simultaneously and is fully independent of our annotation, provides a first step in this direction: it showed that the final boundary under-calls the TIF-seq boundary for fewer than 5% of genes at either end (**Fig. 4**), confirming that over-extension, rather than truncation, is the dominant residual error. A systematic, data-driven re-annotation permitted to shorten over-extended boundaries directly, using such maps to quantify the residual false-positive UTR length discussed above, remains a natural next step. As Nanopore read lengths and direct-RNA chemistry continue to improve, the 5′ resolution that currently limits 5′ UTR calls will tighten, allowing the reference to be refreshed simply by re-running the released pipeline on new data. More broadly, integrating this UTR-focused resource with isoform-discovery efforts such as UNAGI would combine accurate boundary annotation with comprehensive transcript cataloguing, moving the field toward a single, continuously updatable yeast transcriptome reference. Finally, because DRS preserves native RNA base modifications, the same reads can additionally be mined for epitranscriptomic signal alongside boundary annotation (Leger et al., 2021), further increasing the information yield of a single DRS dataset.

### Conclusion

We present a merged *S. cerevisiae* transcriptome reference that extends the Nagalakshmi et al. UTR annotation (Nagalakshmi et al., 2008) using Direct RNA Sequencing, while guaranteeing, by construction and validated independently, that no existing annotation is shortened in the process. The resource fills roughly half of Nagalakshmi’s UTR annotation gaps, improves mapping rate, specificity, and soft-clip burden relative to both the existing reference and naive alternatives on data independent of its construction, and is provided together with its full reproducible pipeline so that it can be applied to new conditions, strains, or sequencing runs as long-read yeast transcriptomics continues to mature. We anticipate this resource will be most immediately useful as a direct substitute reference for existing yeast RNA-seq pipelines, with the underlying pipeline available for groups wishing to extend or refine the annotation further.

## Materials and Methods

### Data accession

Sequencing data used to build the transcriptome reference were generated in (Horvath et al., 2024) and are described in full therein. Briefly, the dataset consists of long-read direct RNA sequencing (DRS) libraries prepared from ribosome-disome-trisome (RDT) and higher polysome (PS) RNA fractions of *S. cerevisiae* (strain BY4741, an S288C background derivative) sampled under non-starved conditions and after 10 minutes of glucose starvation. Libraries were sequenced on an Oxford Nanopore Technologies platform using the SQK-RNA002 chemistry. All raw sequencing data are publicly accessible at the NCBI Sequence Read Archive (SRA) under BioProject PRJNA1022817.

Long read validation data were generated in (Fonzino et al., 2026) and are fully described therein. Briefly, total RNA from *S. cerevisiae* was used for Nanopore libraries preparation with the SQK-RNA002 Direct RNA Sequencing Kit, and was loaded on a single R9.4.1 PromethIon flow cell for sequencing. Raw sequencing data are publicly accessible at the NCBI Sequence Read Archive (SRA) under BioProject SRX28193073.

Short read validation data were generated in (Al Kadi et al., 2020) and are described in full therein. Briefly, *S. cerevisiae* BY4741 strain total RNA was used to generated full-length cDNA using the SMARTer Ultra Low RNA Kit for Illumina Sequencing and sequenced with HiSeq 2500 × 75 bp. The datasets extracted from this study are publicly accessible at DDBJ database (BioProject PRJDB8048) under the accession codes DRR170478-DRR170479.

### Transcriptome reference construction

Two custom transcript (mRNA) references for *S. cerevisiae* were constructed to compare the effect of UTR definition on downstream transcript-level analyses while holding the genome assembly and gene models constant. Both references were built from the SGD S288C genome assembly R64-4-1 (release 20230830 (Cherry et al., 2012)) and a shared canonical single-isoform gene model per locus (gene, CDS and intron features for 6607 genes from the SGD R64-4-1 annotation, reduced to one transcript per gene). The two references differ only in the source of the 5’ and 3’ UTR lengths used to extend each transcript.

#### In-house DRS reference

For the in-house reference, 5’ and 3’ UTR lengths were taken as the longer of the data-driven estimate obtained from our Direct RNA Sequencing data and the Nagalakshmi *et al*. reference annotation, ensuring gene-body coordinates are never more restrictive than the reference annotation. Data-driven boundaries were derived by change-point segmentation of per-position DRS read coverage for each gene (see below).

#### Nagalakshmi reference

For the comparison reference, 5’ and 3’ UTR lengths were taken directly from (Nagalakshmi et al., 2008) genome-wide transcript boundary annotation, in place of the in-house DRS estimates.

For each reference a GFF3 annotation was generated by adding, for every transcript, a 5’UTR and a 3’UTR feature immediately flanking the gene model on the appropriate strand: on the plus strand the 5’UTR spans (gene start - 5’UTR length) to (gene start - 1) and the 3’UTR spans (gene end + 1) to (gene end + 3’UTR length); the assignment is reversed on the minus strand. UTR intervals were clamped to chromosome bounds, and a gene with a zero or sub-2 nt UTR interval received no UTR on that side. Spliced transcript sequences (coding sequence plus UTRs, introns removed) were then extracted from the genome with gffread [v0.12, (Pertea & Pertea, 2020)], with locus clustering and identical-transcript collapsing enabled and the coding extent recorded in each FASTA description (CDS=start-end). Each reference comprises 6,767 transcript sequences.

### In-house DRS reference

Reads were mapped to the *S. cerevisiae* ORF±1000nt reference [SGD S288C assembly R64-4-1] using minimap 2.31 (Li, 2018). Gene-level transcript count profiles were constructed from mapped DRS reads, extending 1,000 nt upstream and downstream of each annotated gene boundary. Genes with fewer than 20 reads at their maximum coverage position were excluded from further analysis. Breakpoint detection was performed gene-by-gene using the Segmentor3IsBack algorithm (Cleynen et al., 2014) with a negative binomial model, which accounts for the overdispersion characteristic of sequencing data [Segmentor(data, model = 3)]. Because the data violate the independence assumptions of the base model, the initial segmentation typically yields a large number of breakpoints. We first remove noisy, closely spaced breakpoints by examining breakpoints within a 40 nt window. For each cluster of nearby breakpoints, we compute the mean coverage on the left and right sides and retain only the breakpoint with the largest absolute difference between these means. We then filter the remaining breakpoints by effect size, keeping only those where the mean coverage difference across the breakpoint exceeds 10% of the gene’s maximum transcript count.

Expressed regions were defined within the 1,000 nt-ORF-1,000 nt window: a segment was called expressed if its mean transcript count exceeded the lower of (i) 5% of the maximum count or (ii) 50 reads. UTR boundaries were extended beyond the 1,000 nt window sequentially if the flanking segment met a stricter threshold of 10% of the maximum (or 50 reads), continuing until coverage dropped below threshold. Final UTR boundaries per gene were taken as the maximum UTR length observed across all sequencing conditions [non-starved / starved] and RNA fractions [PS / RDT]. The overall process is illustrated in **Fig. 1A**.

For each annotated gene, UTR coordinates from our segmentation were compared to the corresponding coordinates in (Nagalakshmi et al., 2008). At each boundary (5′ and 3′ independently), the longer UTR was retained.

### Reproducibility

The complete reconstruction (from segmentation of the data to fasta creation, including chromosome renaming to match annotation seqnames, UTR overlay onto the shared gene-model backbone, and transcript extraction) is provided as a documented pipeline, together with a validation harness. Because the gffread build bundled with cufflinks 2.2.1 is incompatible with the UTR encoding used here, the pipeline builds and uses a standalone gffread 0.12.x, which reproduces the original output exactly. The in-house reference has MD5 f5b10c83168122536a8eed5efe3006cf and the Nagalakshmi reference MD5 fd412507f5fba7d733a26843587f6266. Scripts used to produce the figures in the manuscript, as well as intermerdiary files, are also provided to allow reproducibility from any stage of the pipeline.

### Validation of the transcriptome

#### Comparison of Nagalakshmi *et al*. and long-read UTR annotations

To validate our UTR annotation, we compared the UTR lengths derived from the Nagalakshmi *et al*. short-read sequencing based annotation (Nagalakshmi et al., 2008) with those in our final merged table, on the subset of genes present in both and with non-missing Nagalakshmi estimates. Because our final UTR lengths are defined as the longest UTR observed across all available evidenceincluding the Nagalakshmi annotation itself, a necessary property of a correct merge is that the Nagalakshmi length should never exceed the final length for any given gene. We therefore used this as a primary sanity check, computing the per-gene difference (final − Nagalakshmi) for both 5’ and 3’ UTRs and flagging any gene where this difference was negative.

Beyond this binary check, we examined the overall relationship between the two annotations by computing Spearman and Pearson correlations between Nagalakshmi and final UTR lengths, restricted to genes where both sources reported a non-zero length (this yielded 4,792 and 4,817 comparable genes for 5′ and 3′ respectively). We also compared the distributions of UTR lengths across both annotations and quantified the extent to which the long-read data extended UTR boundaries relative to the Nagalakshmi baseline. All analyses were performed in R using the dplyr, tidyr, and ggplot2 packages and Python using the pysam library.

To benchmark the final annotation against a fully independent transcript-boundary dataset, we additionally compared it with the transcript-isoform sequencing (TIF-seq) map (Gene Expression Omnibus accession GSE39128) (Pelechano et al., 2013), which captures the 5′ and 3′ ends of individual transcripts simultaneously. For each ORF we identified the major covering transcript isoform (mTIF), defined as the most highly read-supported isoform (summed counts across all reported conditions) whose 5′ end lay upstream of the start codon and whose 3′ end lay downstream of the stop codon, and converted its boundaries to per-gene 5′ and 3′ UTR lengths using the ORF coordinates from the gene-model backbone. Because TIF-seq is independent of our reference and plays no part in the longest-UTR merge, boundary differences were treated as signed in both directions and concordance was summarised by Spearman rank correlation and by the fraction of genes agreeing within 20 nt.

### Read alignment

#### Long read DRS

DRS reads were aligned to each reference using minimap2 v2.31. For the genome reference, the splice-aware preset -ax splice was used with the additional flags -uf (forward-strand-only, appropriate for direct RNA sequencing data derived from polyadenylated mRNA) and -k14. For all four transcriptome references (ORF only, ORF ±1000, our annotation, Nagalakshmi), the preset -ax map-ont -k14 -N10 was used. This preset is optimised for Oxford Nanopore reads aligned to a linear (non-spliced) reference. The -N10 flag retains up to 10 secondary alignments per read, enabling quantification of multi-mapping behaviour. All BAMs were coordinate-sorted and indexed with samtools v1.23 (Danecek et al., 2021).

#### Short reads Illumina

For short-read alignment to transcriptome references, STAR v2.7.11b (Dobin et al., 2013) was used. Because STAR requires a pre-built genome index, each transcriptome FASTA was indexed independently using --runMode genomeGenerate with --genomeSAindexNbases 11 and --genomeChrBinNbits 11, parameters scaled to the small size of the yeast transcriptome (approximately 14 Mb for the ORF-only reference, larger for UTR-containing references; the recommended value follows the formula min(14, log2(GenomeLength)/2 − 1)). Reads were aligned with --outSAMattributes NH HI AS NM to retain the NH (number of hits) tag required for multi-mapping quantification. Output was written directly as coordinate-sorted BAM. The two short-read samples were aligned separately and then merged with samtools merge.

Short-read alignment to the genome reference used bwa-mem2 v2.2.1 (Li & Durbin, 2009; Vasimuddin et al., 2019) with default parameters (bwa-mem2 mem). Each sample was aligned separately, sorted, and then merged with samtools merge.

### Alignment quality metrics

The following metrics were computed for every BAM file using samtools v1.23 and custom shell scripts. Unless stated otherwise, secondary (SAM flag 0x100) and supplementary (0x800) alignments were excluded from all counts using samtools view -F 0x900.

#### Mapping rate

For DRS BAMs and the genome-aligned short-read BAM, the mapping rate was computed from samtools flagstat as the fraction of primary reads with at least one alignment. For short-read transcriptome BAMs, STAR does not write unmapped reads to the BAM output by default, causing flagstat to report a spurious 100% mapping rate. The true denominator was therefore computed directly from the input FASTQ files: reads were counted as the number of lines divisible by 4 ( *i.e.*, awk ’NR%4==1’ | wc -l), and the total across both samples (DRR170478 and DRR170479) was used as the denominator for all short-read transcriptome mapping rates. The mapping rate is reported as the number of primary mapped reads in the BAM divided by this FASTQ-derived total.

#### Multi-mapping rate

The metric definition differed by aligner, because minimap2 and STAR use different conventions for recording multiple alignments. For STAR short-read BAMs, the NH auxiliary tag (number of loci the read maps to) is written for every primary alignment. Reads with NH ≥ 2 were counted as multi-mappers using samtools view piped to an awk filter (NH:i: tag parser). The multi-mapping rate is the fraction of all primary reads with NH ≥ 2. For minimap2 DRS BAMs, the NH tag is not written. Multi-mapping was instead quantified from the alignment-per-read distribution: for each read identified by its QNAME, all mapped records in the BAM were counted using a single-pass awk hash table (samtools view -F 0x804 | awk ’{count[$1]++}’), and the frequency distribution of counts was computed. The multi-mapping rate is reported as the fraction of reads with more than one alignment record, derived from this distribution.

#### Per-transcript read counts

Primary alignments for each BAM were extracted with samtools view -F 0x900 -b and re-sorted and indexed, then per-transcript read counts were obtained with samtools idxstats.

#### Per-transcript coverage depth

Mean per-transcript coverage depth and quantised coverage fractions (0×, 1–4×, 5–9×, 10–49×, ≥50×) were computed with mosdepth using the flag --quantize 0:1:5:10:50: (Pedersen & Quinlan, 2018).

#### Antisense alignment rate

For each BAM, primary reads mapping to the reverse strand (SAM flag 0x10) were counted with samtools view -F 0x904 -f 0x10 -c. The antisense rate is the fraction of all primary mapped reads. Because transcriptome FASTA references present all sequences in the sense orientation, a reverse-strand primary alignment represents an antisense read. Note that STAR suppresses antisense alignments by default when aligning to a transcriptome reference; antisense rates for STAR transcriptome BAMs are therefore not interpretable and are reported as not applicable.

### Soft-clip analysis (DRS only)

Soft-clip statistics were computed for DRS transcriptome BAMs using a custom Python script (softclip_metrics.py) implemented with pysam. For each primary alignment, the CIGAR string was parsed to identify leading and trailing soft-clip operations (CIGAR code S, pysam code 4). The lengths of soft-clipped bases at the 5′ and 3′ ends of the read were extracted. For reads mapping to the reverse strand, 5′ and 3′ assignments were swapped to express clipping relative to transcript (rather than read) orientation. Transcripts with fewer than 5 primary alignments were excluded. Per-transcript summary statistics computed include: mean total soft-clip length per read (5′ + 3′ combined); mean clip fraction (total soft-clip bases divided by read length); and asymmetry, defined as the mean per-read difference between 5′ and 3′ soft-clip length.

### Metagene coverage profiles

To compare how each transcriptome reference captures read coverage in the vicinity of ORF boundaries, metagene profiles were computed directly from BAM files using a custom Python script (compute_metagene.py) implemented with pysam, without relying on deepTools.

For each reference, per-base coverage across every transcript sequence in the BAM was computed using pysam.AlignmentFile.count_coverage, restricted to primary alignments (secondary and supplementary reads excluded via a read callback). No additional normalisation was applied beyond per-transcript averaging; the profiles represent mean per base read depth (reads per base) at each position, averaged across all transcripts in the restricted gene set. Transcripts shorter than 200 bp were excluded.

Two metagene profiles were computed per BAM, anchored at the 5′ (TSS) and 3′ (TES) transcript boundaries. For the TSS profile, a vector of length 2 × 1,000 bp was initialised to zero. The first min(1,000, *L*) bases of the per-transcript coverage array were inserted at positions [0, min(1,000, *L*)) relative to the transcript start, where *L* is the transcript length. The preceding positions [−1,000, 0) remained at zero, representing the region upstream of the transcript start boundary, which by definition is absent from all transcriptome references. The TES profile was constructed analogously using the final min(1,000, *L*) coverage values, with positions [0, 1,000) after the transcript end set to zero. The mean and standard error of the mean were then computed position-by-position across all transcripts to yield the metagene profile.

This design allows direct comparison of the four references in a shared coordinate system anchored at ORF boundaries. In the ORF-only reference, no sequence extends beyond the annotated ORF start or end, so coverage is zero at all upstream and downstream positions. In the ORF ±1,000 nt reference, coverage is present in the ±1,000 nt flanks irrespective of UTR length, and may include signal from neighbouring genes given the compact *S. cerevisiae* genome. In the UTR-annotated references (our annotation and Nagalakshmi *et al*.), coverage beyond the ORF boundaries reflects reads originating from the annotated UTR regions, and its shape and extent directly report on UTR length and coverage uniformity.

